# Dual Membrane-spanning Anti-Sigma 2 Controls OMV biogenesis and Colonization Fitness in *Bacteroides thetaiotaomicron*

**DOI:** 10.1101/2025.09.27.678952

**Authors:** Evan J. Pardue, Tengfei Zhong, Nichollas E. Scott, Biswanath Jana, Wandy Beatty, Juan C. Ortiz-Marquez, Mohammed Kaplan, Clay Jackson-Litteken, Mario F. Feldman

## Abstract

*Bacteroides spp.* are Gram-negative, gut commensals that shape the enteric landscape by producing <u>o</u>uter <u>m</u>embrane <u>v</u>esicles (OMVs) that degrade dietary fibers and traffic immunomodulatory biomolecules. Understanding the mechanism behind OMV biogenesis in *Bacteroides spp.* is necessary to determine their role in the gut. Recent studies showed that mutation of <u>D</u>ual <u>M</u>embrane-spanning <u>A</u>nti-sigma factor 1 increased OMV production in *Bacteroides thetaiotaomicron* (*Bt*) by regulating members of its downstream regulon. Additional members of the Dma family have been identified, but very little is known regarding their roles in *Bt*. Here, we investigate the role of <u>D</u>ual <u>M</u>embrane-spanning <u>A</u>nti-sigma factor 2 (Dma2) in controlling OMV biogenesis in *Bt*. We employ biochemical and proteomic analyses to show that mutation of *dma2* increases OMV production in a manner that is dependent on the expression of its cognate sigma factor, *das2*. The precise mechanism by which *dma2* increases OMV biogenesis remains elusive. However, transcriptome analyses revealed that *Δdma2* has decreased expression of select <u>p</u>olysaccharide <u>u</u>tilization loci (PULs) that primarily target host-associated glycans. Follow-up comparative proteomics showed that the PUL repertoire was most impacted in the OMV fraction. *In vitro* growth assessments showed that *Δdma2* exhibits delayed growth in the presence of select host-associated glycans. Colonization studies in mice revealed that *Δdma2* is outcompeted by the wild-type in the gut, which indicates that *dma2* is a key determinant of colonization fitness in *Bt*. Altogether, these findings expand our knowledge of the Dma family’s role in OMV biogenesis and demonstrates their importance in *Bacteroides* physiology.

**Importance:** <u>D</u>ual <u>m</u>embrane-spanning <u>a</u>nti-sigma factors (Dma) are a novel class of regulatory system found solely amongst Bacteroidota. Previous studies have demonstrated the role of Dma1 in vesiculation, but the overall role of the Dma family in *Bacteroides* physiology remains poorly understood. Here, we demonstrate that Dma2 modulates vesiculation and regulates the expression of select <u>p</u>olysaccharide <u>u</u>tilization loci (PULs) that target host-associated glycans. *In vivo* studies revealed that Dma2 is an important fitness determinant when competing against kin bacteria. This work begins characterizing the multifaceted involvement of Dma2 in OMV biogenesis, PUL regulation, and colonization fitness.

## Introduction

The gut microbiota is the consortium of trillions (∼10^12^) of microbes that inhabit the human gastrointestinal tract (Sender et al., 2016; Thursby & Juge, 2017; Zafar & Saier, 2021). This collection of microbes promotes the proper development of the gut epithelium and immune system through various mechanisms (Thursby & Juge, 2017). *Bacteroides spp.* are one of the most abundant genera, making up ∼40% of the bacterial species in the human gut. They help maintain intestinal homeostasis by outcompeting pathogenic microbes, breaking down indigestible dietary fibers, producing short chain fatty acids, and modulating intestinal immunity to reduce inflammation (A. G. Wexler & Goodman, 2017; H. M. Wexler, 2007; Xu et al., 2003; Zafar & Saier, 2021).

*Bacteroides spp.* can stably colonize and successfully compete within the gut due to their ability to utilize a diverse array of dietary polysaccharides and host-associated glycans to promote their growth (A. G. Wexler & Goodman, 2017; Zafar & Saier, 2021). This process is mediated by numerous encoded <u>p</u>olysaccharide <u>u</u>tilization loci (PULs), that can account for ∼20% of the genome in *Bacteroides spp.* PULs are complex nutrient acquisition systems that sense, degrade, and import polysaccharides and other nutrients to be utilized by *Bacteroides spp.* (Hao et al., 2021; Koropatkin et al., 2008; A. G. Wexler & Goodman, 2017; Xu et al., 2003). Each PUL targets a particular class of polysaccharide and are characterized by the presence of SusC and SusD orthologs. SusD-like proteins are surface exposed lipoproteins that bind to polysaccharides, which enables them to be broken down further by surface glycosyl hydrolases, and imported into the periplasm via a SusC-like TonB-dependent <u>o</u>uter <u>m</u>embrane (OM) porin. (Foley et al., 2016; Xu et al., 2003)

Previous mass spectrometry (MS) analyses revealed that <u>o</u>uter <u>m</u>embrane <u>v</u>esicles (OMVs) from *Bacteroides thetaiotaomicron* (*Bt*) and *Bacteroides fragilis* (*Bf*) are preferentially enriched with various surface exposed glycosyl hydrolases, SusD-like proteins, and other proteins typically encoded in PULs (Elhenawy et al., 2014; Sartorio et al., 2023; Valguarnera et al., 2018). OMVs are small, spherical membranous compartments derived from the active blebbing of the <u>o</u>uter <u>m</u>embrane (OM) of Gram-negative bacteria to traffic cellular contents (Sartorio et al., 2021). Due to their glycolytic activity, *Bacteroides* OMVs are viewed as “public goods” because they can degrade various intestinal fibers at a distance and the resulting breakdown products are readily accessible to kin bacteria and other commensal microbes (Rakoff-Nahoum et al., 2014, 2016; Sartorio et al., 2023). Producing OMVs as “public goods” is energetically costly, so *Bacteroides spp.* must tightly control the co-expression of PULs along with OMV biogenesis (Sartorio et al., 2021, 2023). We recently reported evidence for this model demonstrating that *Bt* alters their OMV PUL repertoire to adapt to the extracellular glycan landscape (Sartorio et al., 2023). This phenomenon provides further support for the idea that *Bacteroides* OMVs are important for *Bacteroides spp.* to effectively compete in the gut. Despite their importance, very little is known regarding how OMV biogenesis and regulation occurs. Determining how *Bacteroides* OMVs are produced and regulated is key to understanding the physiology of these microbes and how they function in the gut.

Our recent studies have gained insight into how OMV biogenesis is regulated in *Bt* (Pardue et al., 2024; Sartorio et al., 2023). Briefly, we expressed fluorescent OMV reporters in live cells and employed fluorescence microscopy to visualize OMVs actively blebbing from the OM of live *Bt* cells (Sartorio et al., 2023). By adapting this visualization system, we developed an OMV reporter screen to allow the identification of genes involved in OMV biogenesis and regulation *in vitro* in a high-throughput manner (Pardue et al., 2024). We found that mutation of <u>D</u>ual <u>M</u>embrane-spanning <u>A</u>nti-sigma factor 1 (*Δdma1*) induces OMV production in *Bt* by relieving the repression on its cognate ECF21 family sigma factor, *das1* (Pardue et al., 2024). Dma1 is the first representative of a new class of structurally novel anti-sigma factors that have domains that span from the OM into the cytosol (Pardue et al., 2024). Additional Dma family members, termed Dma2 and Dma3, were identified in *Bt* (Pardue et al., 2024). In this work, we investigate the role of Dma2 in *Bt*. Our findings demonstrate that Dma2 plays dual roles in modulating OMV biogenesis and is an important determinant of colonization fitness by regulating host-glycan targeting PULs.

## Results

### Mutation of Dma2 (BT_1558) increases OMV production

Our preliminary experiments suggested that Dma2 controls OMV biogenesis in a similar manner to Dma1 in *Bt* (Pardue et al., 2024). To begin our analysis, we isolated total membranes (inner and outer membranes; TM) and OMVs from the wild-type (WT), *Δdma2*, and its corresponding complemented strain (*Δdma2_Comp_*), and analyzed their protein profiles via SDS-PAGE followed by Coomassie staining. We found that *Δdma2* exhibited a distorted electrophoretic profile when compared to the WT and *Δdma2_Comp_* **(Fig. 1A)**. The OMV fractions obtained from hypervesiculating strains display irregular SDS-PAGE profiles, due to the increased abundance of lipopolysaccharide (LPS), a key OMV structural component, in these samples. (Pardue et al., 2024). Furthermore, we performed LPS Silver Stains to measure relative amounts of LPS and quantified total protein present in the TM and OMV fractions from the WT, *Δdma2*, and *Δdma2_Comp_* **(Fig. 1B, C)**. Although no differences were observed in the content of LPS and proteins in the TM fractions, the OMV fraction from *Δdma2* contained significantly more LPS and protein when compared to the WT and *Δdma2_Comp_* **(Fig. 1B, C)**. Together, these findings all suggest that mutation of *dma2* causes *Bt* to hypervesiculate. To directly quantify OMV production, we isolated OMVs from the WT, *Δdma2,* and *Δdma2_Comp_* strains, imaged them by transmission <u>e</u>lectron <u>m</u>icroscopy (TEM), and then quantified the number of vesicles present per image. Quantification of OMVs by TEM confirmed that *Δdma2* does indeed increase OMV production compared to the WT. Therefore, the observed phenotypes are not solely due to compositional changes in the contents of the OMV fraction in *Δdma2* **(Fig. 1D)**.

**Figure 1:**
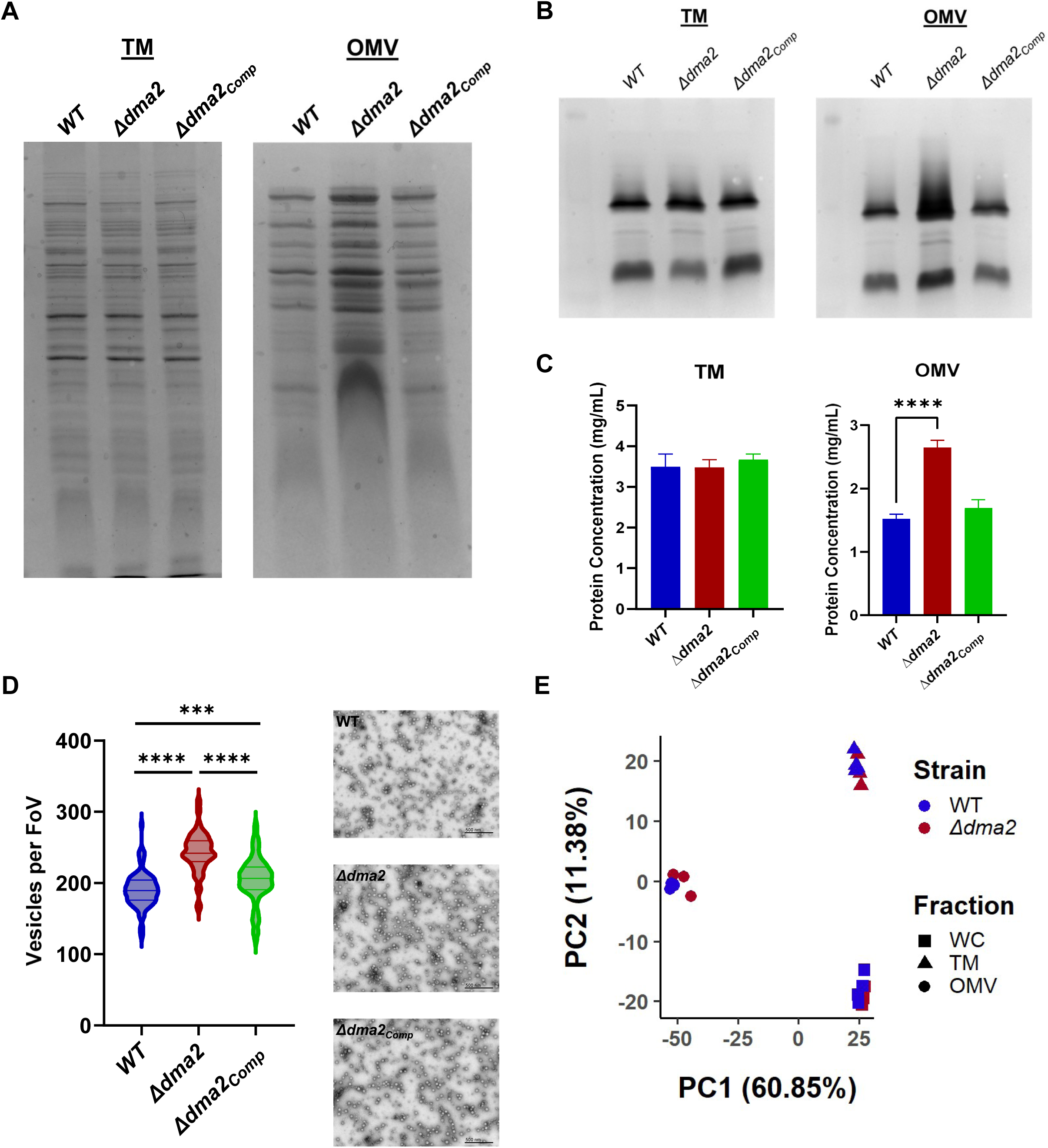
Mutation of Dma2 leads to increased OMV biogenesis in *Bt*. (A) Coomassie Blue stain comparing electrophoretic profiles between TM and OMV fractions from *Bt* WT, Δ*dma2*, and Δ*dma2_Comp_*. Samples were normalized by OD_600_ prior to being run on 10% SDS-PAGE gel. This suggests that deletion of *dma2* induces vesiculation in *Bt*. (B) LPS Silver Stain and (C) Lowry Protein Assay comparing TM and OMV fractions from *Bt* WT, Δ*dma2*, and Δ*dma2_Comp_*. These show that Δ*dma2* contains more LPS and proteins in its OMV fraction, which is consistent with increased OMV production. (D) Transmission electron microscopy (TEM) confirms that *Δdma2* produces significantly more OMVs than the WT. <u>Left:</u> Results of quantification of 90 TEM images of OMVs from the OMV fraction from each strain (FoV: Field of View). <u>Right</u>: Representative TEM images of OMVs from each strain. Three biological replicates of aliquots from the OMV fraction of each strain were fixed onto grids in triplicate (in materials and methods). Ten random images were taken from each grid (n=90 per strain) and OMVs were counted manually. (E) Principal component analysis of WC, TM, and OMV proteomic data from *Bt* WT and Δ*dma2*. This demonstrates that the overall composition of each cellular fraction are similar between the two strains.

Previous studies have demonstrated that *Bacteroides* OMVs contain select protein cargo that consists primarily of surface exposed lipoproteins derived from <u>p</u>olysaccharide <u>u</u>tilization loci (PULs) (Elhenawy et al., 2014; Sartorio et al., 2023; Valguarnera et al., 2018). To ensure that the increase in vesiculation in *Δdma2* is not caused by cell lysis, we performed comparative proteomic analyses of whole cells (WC), TMs, and OMV from the WT and *Δdma2*. The resulting principal component analysis (PCA) shows that each fraction from the WT and *Δdma2* contain similar protein composition **(Fig. 1E)**. Vesicles generated by lysis usually carry ribosomal proteins and other cytoplasmic components. However, since the proper OMV cargo selection is maintained in *Δdma2*, we can rule out that the increased OMV production observed in *Δdma2* is due to membrane instability and cell lysis.

Next, we performed cryo-electron tomography (cryoET) on the WT and *Δdma2* to assess the morphology of *Bt* OMVs. The size and electron density of OMVs was not impacted in *Δdma2* **(Fig. 2A, B**). However, we found that *Δdma2* exhibited a higher proportion of coccoid cells (48.3%) when compared to the WT (24.4%) **(Fig. 2C-F)**. This suggests a potential link between OMV production and cell shape maintenance.

**Figure 2:**
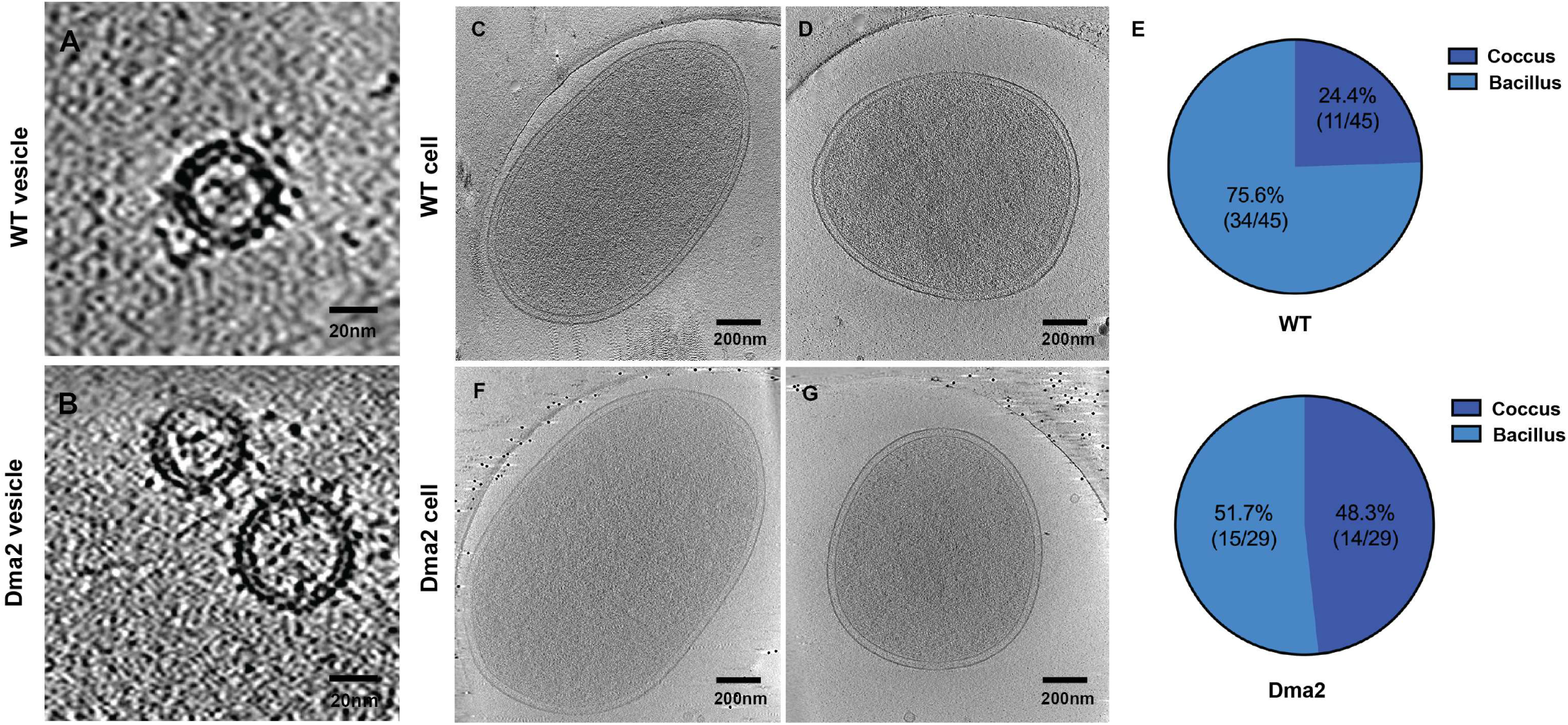
Mutation of Dma2 causes morphological changes in *Bt* cells and OMVs. Representative cryo-electron tomographs of OMVs from (A) *Bt* WT and (B) *Δdma2.* Representative cryo-electron tomographs of rod-shaped and coccoid, and their corresponding ratios from (C-E) *Bt* WT and (F-H) *Δdma2*. This shows that mutation of *dma2* increases the number of coccoid cells present in *Bt*.

### Das2 (BT_1559) is required to induce OMV production in *Δdma2*

Dma2, like other Dma family members, exhibits a unique domain organization, consisting of (1) an N-terminal anti-sigma binding domain, (2) a transmembrane helix, (3) a long, intrinsically disordered tether-like region, and (4) a C-terminal β-barrel domain **(Fig. 3A)**. We showed previously that Dma1 modulates OMV biogenesis by directly controlling the activity of its cognate sigma factor, Das1 (Pardue et al., 2024). Dma2 is encoded in a five gene operon containing a putative ECF21 family sigma factor (*das2*; BT_1559) and three additional genes (BT_1555, BT_1556, and BT_1557) **(Fig. 3B)**. ECF21 family sigma factors are found solely amongst Bacteroidota and are encoded adjacent to Dma family members (Ndamukong et al., 2013; Pardue et al., 2024; Staroń et al., 2009). This strongly suggests that Dma2 and Das2 form a sigma/anti-sigma pair. Because of this, we hypothesized that the hypervesiculation observed in *Δdma2* is due to the liberation of Das2 (**Fig. 3C)**. To confirm whether Das2 is required to induce OMV biogenesis in *Bt*, we generated clean *das2* deletion mutants in the WT and *Δdma2* background. Growth curves were performed with these strains and showed that there is no significant fitness defect in rich media **(Fig. S1)**. Next, we isolated OMVs from the WT, *Δdma2, Δdas2, Δdma2-das2* and *Δdma2_Comp_* and compared the electrophoretic profiles, as a proxy for OMV biogenesis, by SDS-PAGE followed by Coomassie staining. While we observed no phenotype in *Δdas2*, we found that *Δdma2-das2* displayed a WT electrophoretic profile **(Fig. 3D)**. This confirms that the increased OMV production observed in *Δdma2* requires the activity of its sigma factor, *das2*.

**Figure 3:**
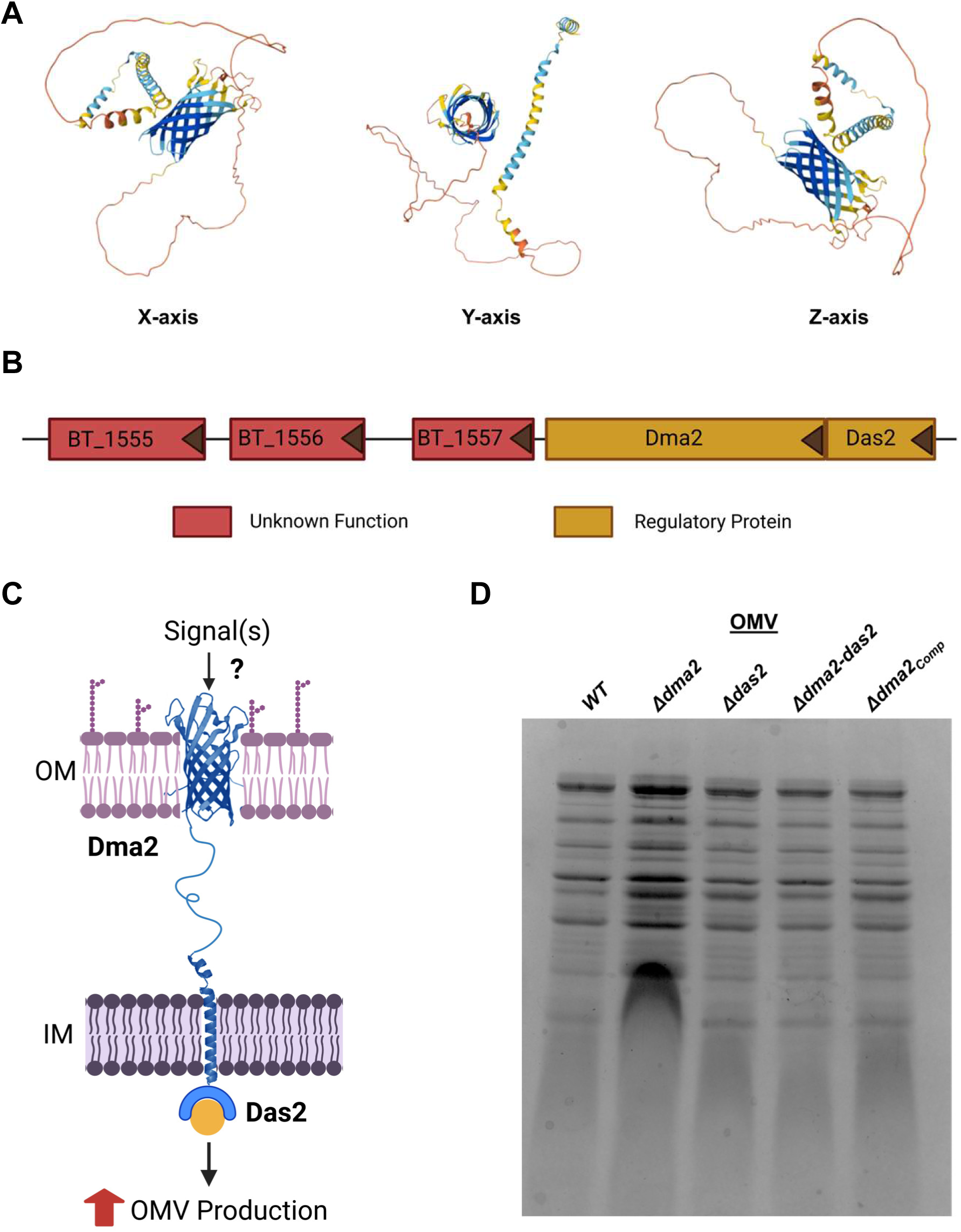
ECF21 family sigma factor, *das2,* is required to induce OMV biogenesis in *Bt*. (A) AlphaFold structural predictions of Dma2. (Varadi et al., 2022)(B) Schematic and putative functions of each gene present in the *dma2* operon (Created with BioRender.com). (C) Proposed model of how Dma2 induces OMV production in *Bt* by controlling the activity of Das2 (Created with BioRender.com) (D) Coomassie Blue stain comparing electrophoretic profiles of OMV fractions from *Bt* WT, Δ*dma2*, Δ*das2*, Δ*das2-dma2*, and Δ*dma2*_Comp_. Samples were normalized by OD_600_ values and run on 10% SDS-PAGE gel. This confirms that deletion of *das2* in the *Δdma2* background restores WT levels of vesiculation.

### Dma2 controls OMV biogenesis in a manner that is distinct from that of Dma1

To investigate how Dma2 controls vesiculation, we compared the transcriptome of *Δdma2* to that of the WT. Our analysis revealed that the most differentially regulated genes belonged to two main categories: (1) genes encoded in the same operon as *dma2* and (2) genes that are part of PULs **(Fig. 4A; Dataset S1, S2)**.

**Figure 4:**
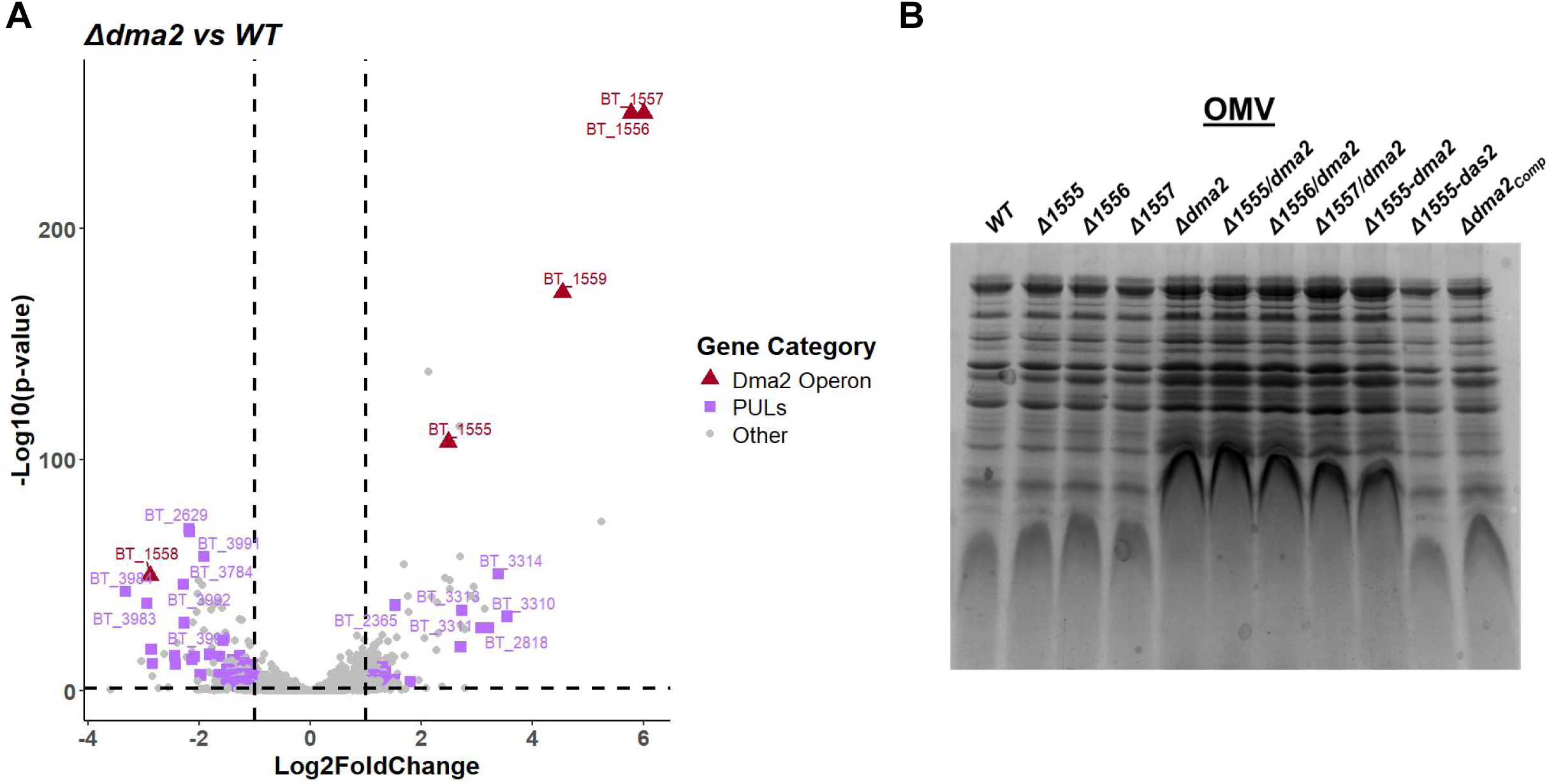
Dma2 primarily regulates its own operon and PULs targeting host-associated glycans in *Bt.* (A) Volcano plot representations of transcriptome data comparing *Bt* WT and *Δdma2* **(Dataset S1, S2)**. (B) Coomassie Blue stain of OMV fractions isolated from *Bt* WT and strains containing deletions in the downstream genes of the *dma2* operon. Samples were normalized by OD_600_ and run on 10% SDS-PAGE. This experiment shows that the *dma2* operon downstream genes are not responsible for the induction of OMV production observed in *Δdma2*.

In *Δdma2,* BT_1555 (Log_2_FC: 2.48), BT_1556 (Log_2_FC: 5.78), BT_1557 (Log_2_FC: 6.00), and *das2* (BT_1559) (Log2FC: 4.54) were the most upregulated genes **(Fig. 3B**; **Fig. 4A; Table S2; Dataset S1)**. No studies have attributed functions to these gene; however, Foldseek predicts that BT_1555 is structurally similarity to enoyl-acyl carrier protein (ACP) reductases and nitronate monooxygenases, while BT_1556 and BT_1557 are annotated as DUF4858- and DUF4943 domain containing proteins, respectively (van Kempen et al., 2024). To test whether these genes impact OMV production, we generated mutant strains in each of these genes in the *Δdma2* background. OMV analysis revealed that deletion of none of the downstream genes in the *dma2* operon have an impact on OMV biogenesis **(Fig. 4B)**.

We previously showed that *Δdma1* induces the expression of *nigD1* along with other select genes in *Bt* (Pardue et al., 2024). NigD1 belongs to a class of proteins called NigD-like proteins that are found solely amongst Bacteroidota, and it is required to increase OMV production in *Δdma1* (Pardue et al., 2024). Since Dma1 and Dma2 both belong to the same family and increase OMV biogenesis, we hypothesized that mutation of *dma2* could also induce OMV production by increasing the expression of *nigD1*. However, our RNAseq analysis revealed that the expression of *nigD1* (Log_2_FC: -0.79) was slightly downregulated in Δ*dma2*, which counters this idea **(Table S2; Dataset S1)**.

Lastly, to determine whether there are shared genes regulated by both *dma1* and *dma2* that could provide additional insight regarding how they modulate OMV biogenesis, we compared our previous transcriptome data from *Δdma1* to that collected from *Δdma2*. We found that genes encoding a putative type V pilus (Bt_2655-2660) (Mihajlovic Jovana et al., 2019), an orphan ECF type sigma factor (BT_2569), the *dma3* locus (BT_2778-2779) (Pardue et al., 2024), and many hypothetical proteins are upregulated, while components from PUL36 (α-mannan/host N-glycans), PUL52 (unknown), PUL67 (mucin-O-glycans), and PUL68 (α-mannan/host N-glycans) (Cuskin et al., 2015; Martens et al., 2008), S-layer proteins (BT_1926-1927) (Taketani Mao et al., 2015), and glycine betaine/L-proline transport system permeases (BT_1750-1751) (Yaung et al., 2015) are downregulated in both *Δdma1* and *Δdma2* **(Table S3; Dataset S3, S4)**. Many of the shared genes identified here are not likely to be implicated in OMV biogenesis, but the *dma3* locus is of significant interest. Since the *dma3* locus is upregulated in both strains, this suggests that there is potentially crosstalk occurring between members of the Dma family. To determine whether *dma3* plays a role in inducing OMV biogenesis, we generated clean deletion mutants in *dma3* in the WT, *Δdma1*, and *Δdma2* backgrounds prior to isolating the OMV fraction and visualizing the electrophoretic profile by SDS-PAGE followed by Coomassie staining. However, mutation of *dma3* does not revert the distortion in the electrophoretic profiles observed in *Δdma1* and *Δdma2*. This indicates that *dma3* activity is not required for these strains to hypervesiculate **(Fig. S2)**. Additional studies are required to determine the role of Dma3 in *Bt*. Altogether, our findings support the conclusion that *Δdma1* and *Δdma2* induce OMV production through distinct regulatory cascades.

### Absence of Dma2 results in altered OMV cargo selection

PULs are complex, nutrient acquisition systems encoded by Bacteroidota that enable them to utilize a wide array of dietary-, microbial-, and host-derived glycans (Hao et al., 2021; Koropatkin et al., 2008; Martens et al., 2008; A. G. Wexler & Goodman, 2017). The transcriptomic data from *Δdma2* revealed that genes in select PULs are differentially regulated. We found that PUL36 (α-mannan/host N-glycans), PUL52 (unknown substrate), PUL68 (α-mannan/host N-glycans), and PUL72 (mucin-O-glycans/high mannose mammalian N-glycan) are the most repressed PULs when compared to the WT. On the other hand, PUL38 (mucin-O-glycans) and PUL56 (1,6-β-Glucan) were induced in *Δdma2* when compared to the WT **(Fig. 4A**; **Table 1; Dataset S1, S2)**.

**Table 1:**
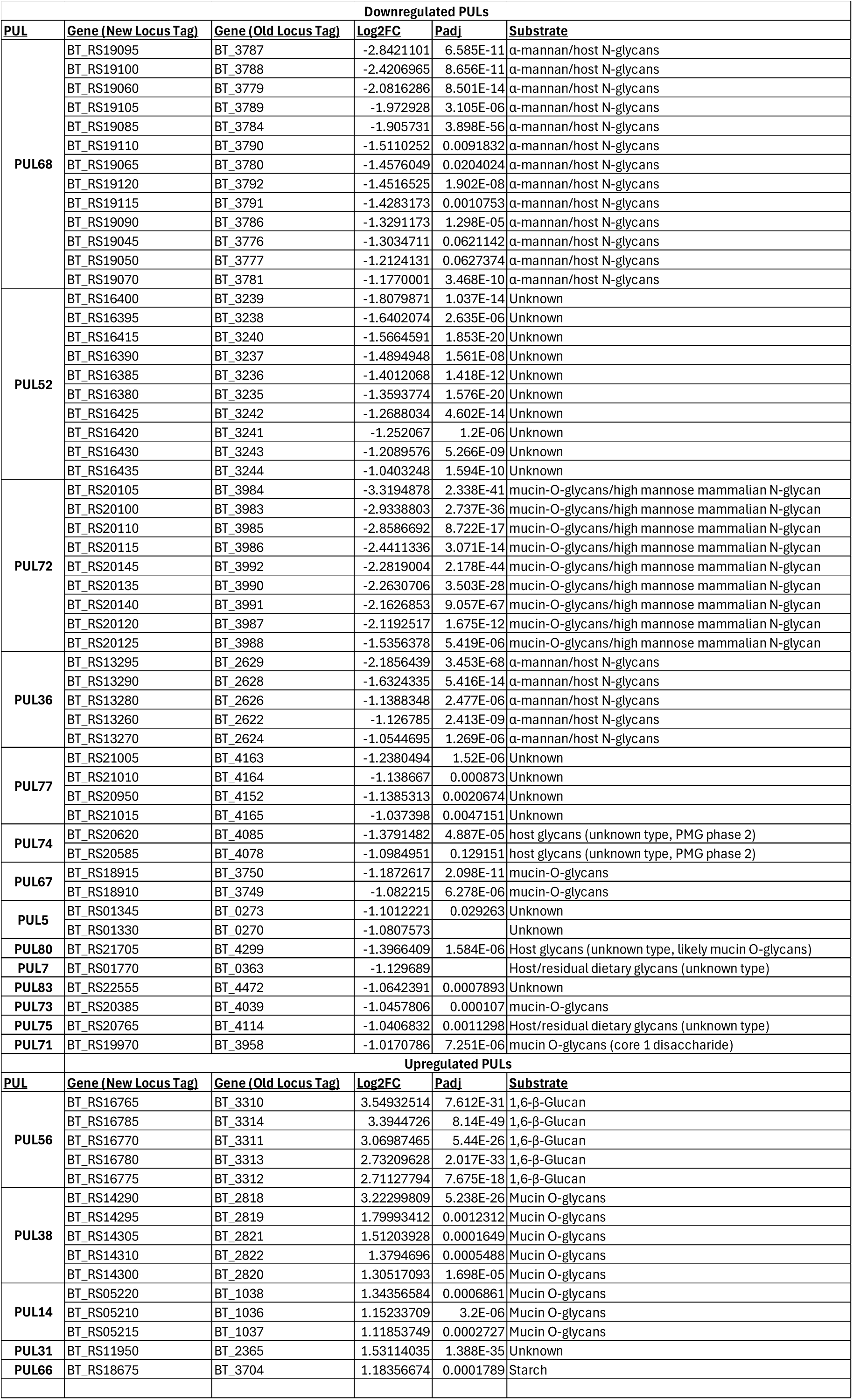
Dysregulation of the PUL repertoire occurs in Δdma2.

Since Dma2 is an anti-sigma factor that functions by controlling the activity of Das2, we propose that the selective dysregulation of PULs observed in *Δdma2* is caused by *das2* activity. The WT bacteria express basal levels of Das2, so to better assess the *das2*-specific phenotypes, we conducted additional transcriptomic analyses comparing *Δdma2* (ON-state) to *Δdma2-das2* (OFF-state). Like the previous RNA-seq, the downstream genes in the *dma2* operon were the most induced, while the expression of various PULs were altered **(Table S4; Dataset S5)**. In this new data set, additional genes related to capsule biosynthesis were also differentially regulated, which we postulate is a consequence of phase variation, a common phenomenon in Bacteroides **(Dataset S5)** (Hsieh et al., 2021; Porter et al., 2020). Remarkably, PUL1 (unknown), PUL31 (unknown), PUL38 (mucin-O-glycans), PUL56 (1,6-β-Glucan), and PUL69 (α-mannan/host N-glycans), were significantly induced, while PUL22 (levan/fructooligosaccharides), PUL36 (α-mannan/host N-glycans), PUL37 (ribose/ribonucleosides), PUL68 (α-mannan/host N-glycans), PUL72 (mucin-O-glycans), PUL84 (mucin-O-glycans) were significantly repressed in *Δdma2* **(Fig. 5A; Table S5; Dataset S5)**. A majority of the differentially expressed PULs target host-associated glycans (Martens et al., 2008).

**Figure 5:**
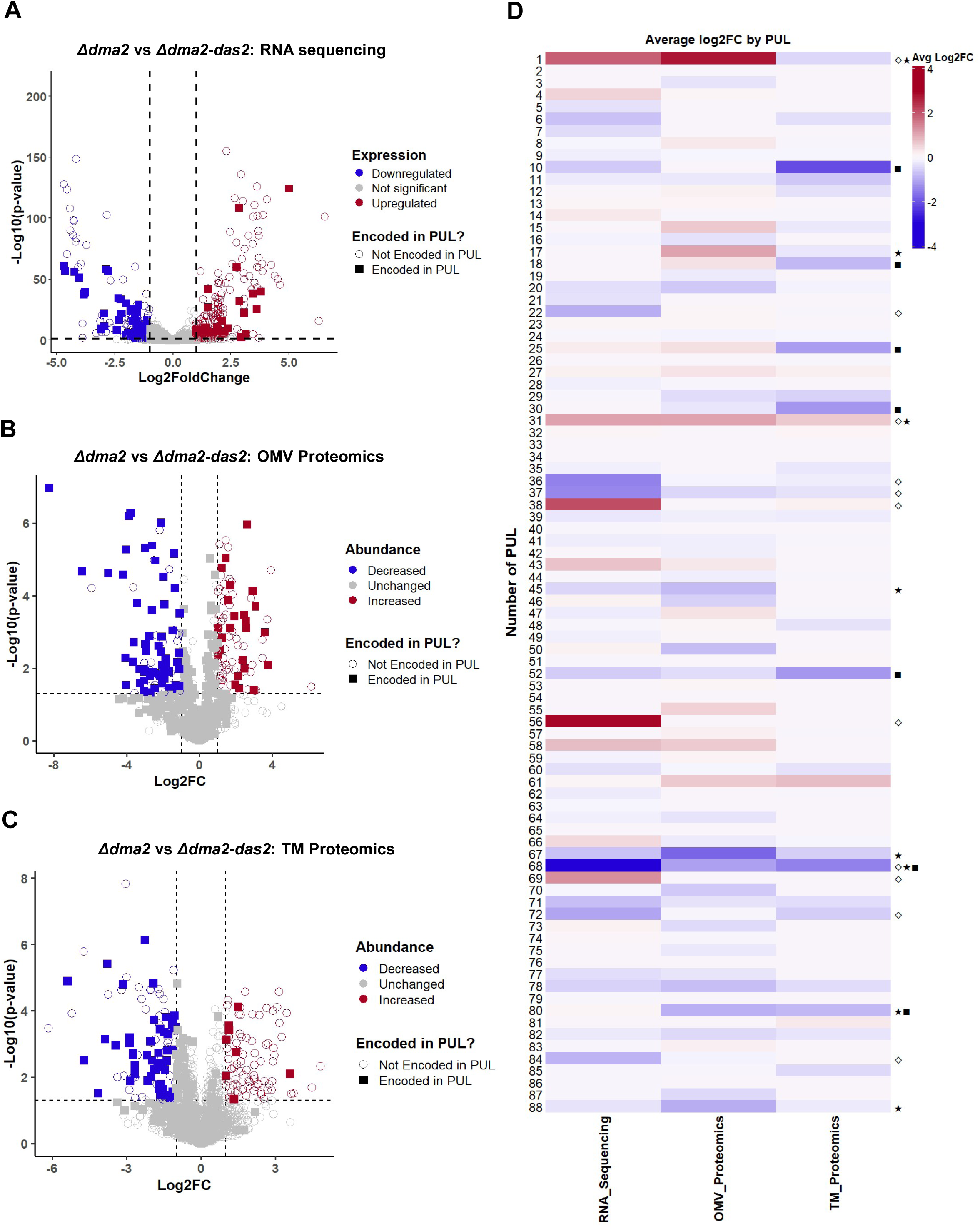
Select PULs targeting host-associated glycans are downregulated in *Δdma2.* Volcano plot representations of (A) RNAseq, (B) WC, (C) TM, and (D) OMV proteomic data comparing *Bt Δdma2* and *Δdma2-das2* **(Dataset S5-S7)**. These analyses reveal that select PULs primarily targeting host-associated glycans are significantly repressed in *Δdma2.* These changes are most prevalent in the OMV fraction. (E) Heatmap comparing the differential expression (Average log2FC) of each PUL present in *Bt* from *Bt Δdma2* and *Δdma2-das2* transcriptome and proteome datasets. Average log2FC was calculated by filtering out each gene found to be encoded within a particular PUL and averaging the significant log2FC values (log2FC ≥ |±1| and p-value < 0.05). Non-significant log2FC values were included when calculating the Average log2FC for each PUL, but these were assigned the arbitrary value of 0. This analysis enables us to distinguish whether entire PULs are altered in *Δdma2* or if only specific components of certain PULs are changed. PULs that are significantly altered are indicated by the following symbols: “◊” for RNA sequencing, “▪” for TM proteomics, and “★” for OMV proteomics.

To evaluate whether the modulation of PUL expression observed in *Δdma2* affects OMV cargo, we performed comparative proteomic analyses on OMVs isolated from *Δdma2* and *Δdma2-das2*. In our analysis, we also included total membranes (inner and outer; TM). In *Δdma2* OMVs, PULs comprised ∼50% of the total proteins found to be significantly altered. On the contrary, the TM fraction displayed less changes **(Fig. S3; Dataset S6, S7).** Our analyses revealed that PULs are primarily repressed at the protein level in these subcellular fractions of *Δdma2*, while very few are induced **(Fig. 5B, C; Table S6; Dataset S6, S7)**. We found that of the PULs shown to be differentially expressed in the RNA sequencing, only PUL1, PUL31, and PUL68 were also altered in the OMV fraction **(Fig. 5D; Dataset S5-S7)**. In addition, PUL17 (host glycans), PUL45 (host glycans), PUL67 (mucin O-glycans), PUL80 (host glycans), and PUL88 (unknown) are altered in the OMV fraction, while PUL10 (unknown), PUL18 (unknown), PUL25 (galactooligosaccharides), PUL30 (mucin), PUL52 (unknown), and PUL80 (host glycans) are changed in the TM fractions, even though these PULs were not differentially expressed at the transcriptional level **(Fig. 5D; Dataset S5-S7)**. Previous studies have shown that *Bt* selectively tailors its OMV cargo based on the extracellular nutrient landscape (Sartorio et al., 2023). However, our findings are the first to implicate the Dma family in shaping OMV protein cargo through the modulation of PULs in *Bt*.

### *Δdma2* exhibits delayed growth on select host-associated glycans *in vitro*

Since *Δdma2* represses the activity of many PULs that primarily target host and microbial glycans **(Fig. 5D; Table S5, S6; Dataset S5-S7;)**, we hypothesized that *Δdma2* would exhibit stunted growth compared to the WT when grown in minimal media supplemented with these types of glycans as a sole carbon source. PUL68 is important for growth in the presence of yeast α-mannan and was the only PUL shown to be significantly repressed in *Δdma2* for each of our comparative analyses **(Fig. 5D)**. However, we found that *Δdma2* does not exhibit delayed growth *in vitro* compared to the WT and complemented strain when grown in the presence of yeast α-mannan **(Fig. 6A)**. This suggests that either (1) the repression of PUL68 observed in *Δdma2* is not sufficient to delay the expression of this PUL when α-mannan is present, (2) PUL36 and PUL69, which also target yeast α-mannan, may have adequate expression to compensate for the repressed PUL68, or (3) the yeast α-mannan utilized for this experiment may not be the proper substrate (Cuskin et al., 2015).

**Figure 6:**
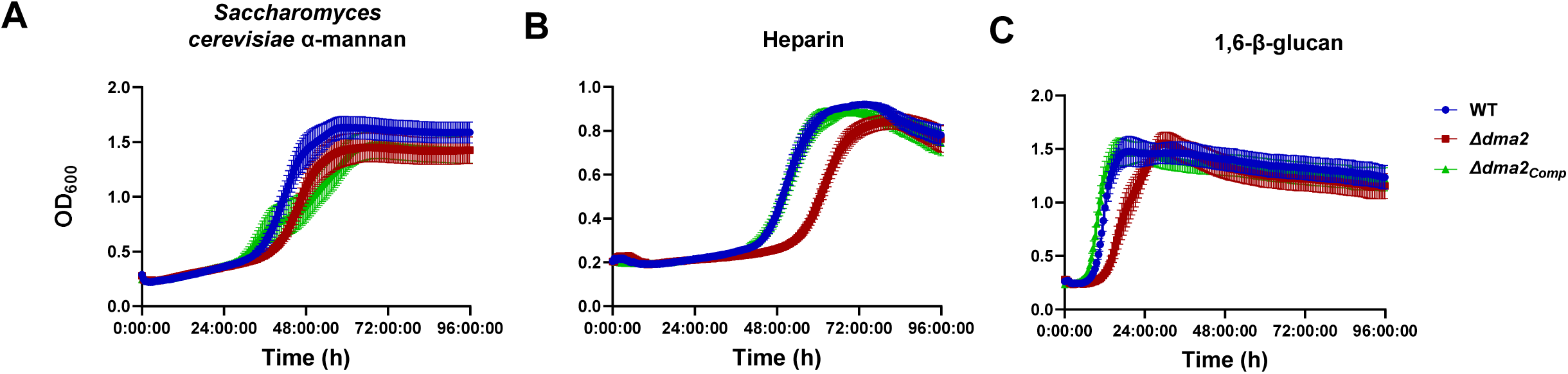
*Δdma2* exhibits delayed growth on select host- and microbially-derived glycans. Growth curves showing the growth of *Bt* WT, *Δdma2*, and *Δdma2_Comp_* in the presence of minimal media containing (A) α-mannan from *Saccharomyces cerevisiae*, (B) heparin, and (C) 1,6-β-glucan. Growth curves were generated from the results of three independent experiments each including four technical replicates from each strain.

To determine whether the dysregulation of PULs in *Δdma2* impacts their growth on different sole carbon sources *in vitro*, we tested a panel of monosaccharides (glucose, fructose, galactose, arabinose, mannose, rhamnose, and xylose), plant polysaccharides (amylopectin, gum arabic, levan, pectin, and rhamnogalacturonan), and host- and microbially-derived glycans (hyaluronan, heparin, 1,6-β-glucan, and porcine mucin type II/III). We found that *Δdma2* had growth comparable to the WT when monosaccharides or plant polysaccharides were the sole carbon source **(Fig. S4)**. However, *Δdma2* exhibited delayed growth on two nutrients: heparin, which is present in the intestinal mucosa and secreted by mast cells in the gut, and 1,6-β-glucan, which is primarily found in yeast and fungal cell walls **(Fig. 6B, C)** (Cartmell et al., 2017; Temple et al., 2017). Our findings confirm that *Δdma2* is less able to utilize select host-associated glycans *in vitro*.

### Dma2 is an important determinant of *in vivo* fitness in *Bt*

PULs that target host-associated glycans are important for *Bt* to stably colonize the human gut (Martens et al., 2008). Since *Δdma2* displayed delayed growth on heparin and 1,6-β-glucan **(Fig. 6B, C)**, we hypothesized that *Δdma2* may be defective in colonizing *in vivo*. To test this, we treated C57/BL6 mice with an antibiotic cocktail for 7 days prior to colonizing with either *Bt* WT or *Δdma2* by oral gavage and measuring CFUs in feces in intervals for 14 days **(Fig. 7A)**. Mono-colonization experiments revealed that *Δdma2* colonized to the same degree as the WT in our antibiotic-treated mouse model **(Fig. 7B, C)**. On the other hand, co-colonization with the WT and *Δdma2* strains revealed that *Δdma2* initially colonizes at higher levels than the WT, but after 5 days post oral gavage, *Δdma2* rapidly declines in abundance, while the WT remains stable in the population **(Fig. 7D, E)**. To ensure that the fitness defect in *Δdma2* is not due to the overexpression of downstream genes in the *dma2* operon, we also co-colonized with the WT and *Δ1555-dma2,* which lacks *dma2* and the rest of the operon besides *das2*. We confirm that *Δ1555-dma2* is still outcompeted by the WT, but unlike *Δdma2*, which initially colonizes better than the WT, *Δ1555-dma2* starts to decline in abundance immediately before appearing to stabilize **(Fig. S5A, B)**. Altogether, these findings demonstrate that Dma2 is an important determinant of *in vivo* fitness in *Bt*.

**Figure 7:**
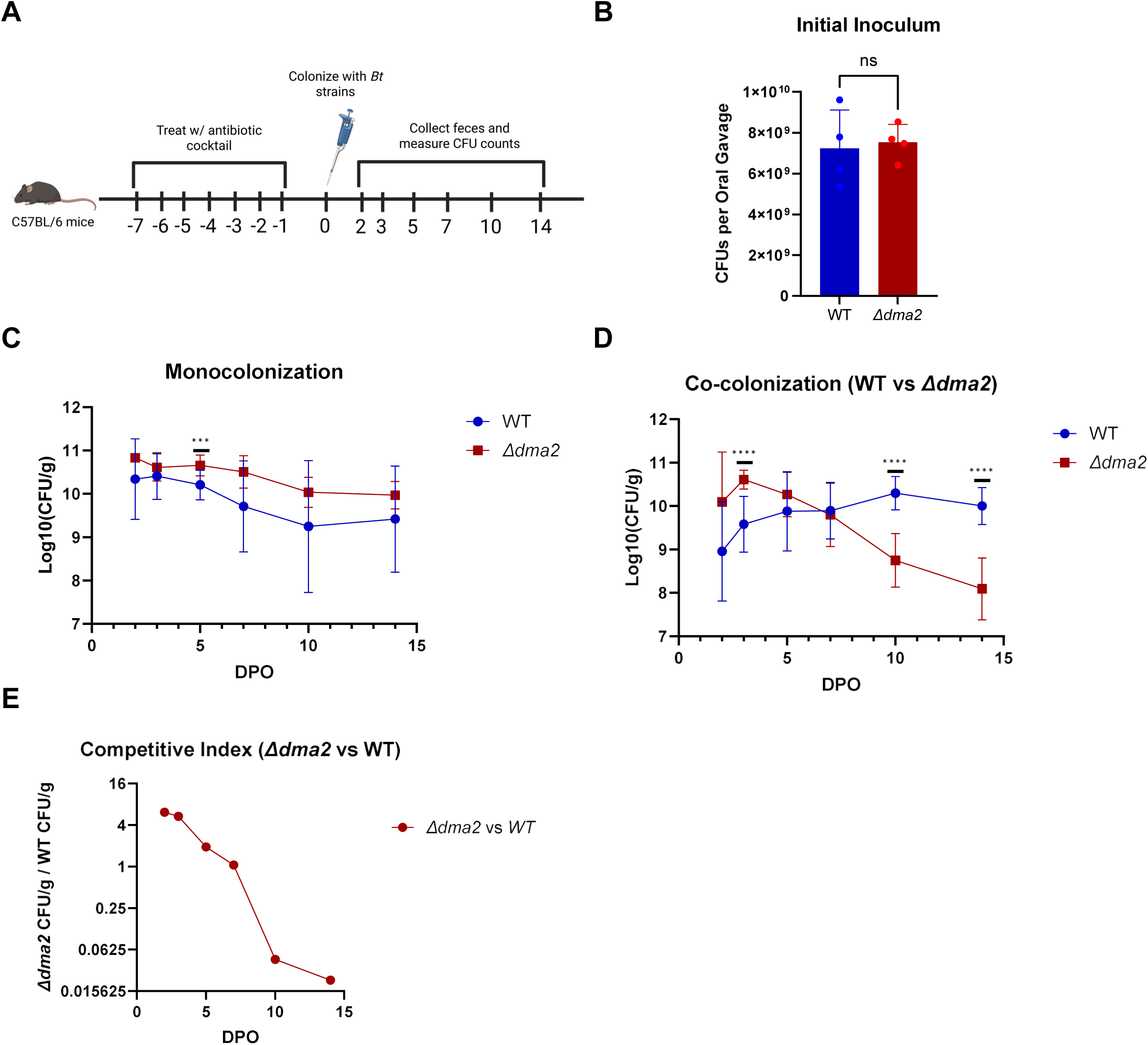
Dma2 is an important colonization factor in *Bt*. (A) Schematic outlining *in vivo* colonization of antibiotic-treated mice experiments (Created with BioRender.com). (B) Monocolonization studies comparing *Bt* WT, *Δdma2* and *Δdma2-das2*. This experiment shows that *Bt* WT, *Δdma2* and *Δdma2-das2* colonize to comparable levels *in vivo*. (C, D) Co-colonization experiment comparing *Bt* WT and *Δdma2*. This shows that *Δdma2* is unable to maintain stable colonization when the WT is present. (E, F) Co-colonization experiment comparing *Bt* WT and *Δdma2-das2*. This reveals that *Δdma2-das2* colonizes at higher levels than the WT throughout the duration of the experiment. Points on the graph represent the mean and standard deviation of data collected from three independent experiments containing four mice each per condition. Significance threshold corresponds to: (*) p-value = 0.05, (**) p-value = 0.01, (***) p-value = 0.001, (****) p-value = 0.0001.

## Discussion

The Dma family is a novel class anti-sigma factors found solely amongst Bacteroidota (Evans et al., 2022; Ndamukong et al., 2013; Pardue et al., 2024). To date, members of the Dma family are known to play a role in OMV biogenesis, but we still lack a complete understanding of their role in these microbes. In this study, we showed that deletion of *dma2* in *Bt* results in a significant increase in vesiculation. Dma2-mediated hypervesiculation requires the activity of its cognate sigma factor, Das2. In addition, transcriptome and proteomic analyses show that Dma2 and Das2 are required for proper OMV cargo selection. Lastly, we demonstrate through *in vivo* studies that Dma2 is a key determinant of colonization fitness.

Dma1 and Das1 induce OMV production by increasing the expression of *nigD1* (Pardue et al., 2024). On the contrary, we demonstrated that Dma2 and Das2 increase OMV production, but do not increase the expression of NigD1 **(Table S2; Dataset S1)**. This suggests that Dma1 and Dma2 regulate OMV biogenesis through distinct regulatory cascades. Even so, the role of NigD1, and other NigD-like proteins, in *Bt* is currently unknown. Future studies are required to understand the precise mechanism by which Dma2 controls OMV production.

In *Δdma2*, OMV size and structure is not affected **(Fig. 2A, B)**, but the cells displayed a higher abundance of coccoid cells **(Fig. 2C-H)**. To produce OMVs, cells must properly coordinate the production and trafficking of LPS, other membrane lipids, and OM proteins. This process is likely energetically costly to the bacteria because the secreted cellular contents must be replenished. Interestingly, our findings could indicate that the induction of OMV biogenesis in *Δdma2* impacts the structure of the cell envelope due to an imbalance between the production and export of OM contents.

We previously showed that when *Bt* is grown in the presence of different polysaccharides, they package their OMVs with enzymes required to degrade them (Sartorio et al., 2023). This established a direct link between PUL induction and OMV cargo selection. Interestingly, mucin is a special case where *Bt* induces the requisite PULs, but these components are retained at the cell surface, instead of localizing to OMVs (Sartorio et al., 2023). Here, we demonstrate that *Δdma2* primarily alters the expression of various PULs, but many of these transcriptional changes are not necessarily conserved at the protein level **(Fig. 5A-D)**. This makes Dma2 the first gene shown to impact OMV cargo selection. Most significantly, some of these changes in OMV cargo occur independent of transcriptional regulation, which indicates that additional regulatory features are at play that impact OMV cargo selection. It remains a possibility that Dma2 functions to preclude specific components from mucin-targeting PULs from reaching OMVs. This is physiologically important because *Bt* seeks to avoid degrading the host’s mucus lining in an uncontrolled fashion because this has been shown to cause inflammation in certain dietary contexts (Desai et al., 2016; Sartorio et al., 2023).

Comparisons of our transcriptome analyses between *Δdma1* and *Δdma2* revealed that *dma3* is significantly induced in both cases. Very little is known about Dma3, except that it is the most structurally unique member of the Dma family because it is not encoded adjacent to a cognate ECF21 family sigma factor, instead Dma3 encodes a domain that functions as its cognate sigma factor at the N-terminus of the protein (Pardue et al., 2024). We showed that Dma3 does not impact OMV production in *Δdma1* and *Δdma2* **(Fig. S2)**. Comparative transcriptome analyses revealed that *Δdma1* and *Δdma2* both impact the expression of select PULs **(Table S3; Dataset S3, S4)**, so it remains a possibility that Dma3 could play a role in this process. Overall, understanding the role of the Dma family in regulating the PUL repertoire is of significant interest.

In *Bacteroides spp.*, PULs are primarily controlled by SusR-like regulators (Cho Kyu Hong et al., 2001; Foley et al., 2016), hybrid two component systems (HTCS) (Gao Rong et al., 2024; Lee et al., 2022; Lynch & Sonnenburg, 2012; Schwalm III et al., 2017), and ECF type sigma/anti-sigma factor systems (Martens et al., 2008). These systems enable *Bt* to maintain a strict glycan hierarchy where monosaccharides and plant polysaccharides are preferred over host mucosal glycans (Martens et al., 2008; Pudlo Nicholas A. et al., 2015; Rogers et al., 2013). PULs that target mucin and other host glycans are known to disproportionately be associated with ECF type sigma/anti-sigma factors. However, to the best of our knowledge, this class of regulators has yet to be shown to be important for maintaining the glycan hierarchy in *Bacteroides spp.* (Martens et al., 2008). Our data suggests that Dma2 is involved in this process by downregulating select PULs that target host-associated glycans **(Fig. 5A, Table S2, S4; Dataset S1, S2, S5)**. Because of this, we propose that Dma2 can sense the presence of a currently unknown glycan at the OM surface and then releases Das2 to directly repress the activity of PULs from lower priority host glycan targeting PULs.

Our experiments provide preliminary support for this model. In **Figure 6**, we demonstrate that *Δdma2* exhibits delayed growth when heparin or 1,6-β-glucan is the sole carbon source. In *Bt*, heparin has been considered a high priority glycan because PUL85, which targets heparin, is not significantly repressed when monosaccharides are present (Rogers et al., 2013). Our proteomic analyses showed that BT_4659 (SusD-like) and BT_4660 (SusC-like) from PUL85 are repressed in *Δdma2* **(Table S6; Dataset S6, S7)**, which suggests that Dma2 could sense a higher priority polysaccharide. In tandem, Das2 could modulate the activity of these lower priority PULs by inhibiting the activity of their native regulatory systems (Martens et al., 2008). Future studies will aim to elucidate the substrates that are sensed by Dma2 and decouple whether Das2 functions by directly or indirectly repressing the activity of host glycan targeting PULs in this context.

Tightly controlling the expression of PULs is key for *Bacteroides spp.* to adapt to and thrive in the human gut (Martens et al., 2008). Our experiments revealed that lack of *dma2* drastically impacts the ability of *Bt* to maintain stable colonization when there is competition from other kin bacteria **(Fig. 7D, E)**. We hypothesize that the dysregulation of PULs and OMV cargo selection in *Δdma2* causes the observed fitness defect, but many questions remain. The *dma2* operon downstream genes (BT_1555-BT_1557) represent the most upregulated genes in *Δdma2.* Our findings demonstrate that these genes are not involved in OMV biogenesis but could be important for the initial colonization fitness of *Δdma2 (***Fig. S5)**. The function of these genes has not been experimentally confirmed, so elucidating the exact function of these genes is required to understand their role *in vivo*.

The Dma2 operon is only encoded in *Bt* and closely related *Bacteroides spp* (Pardue et al., 2024). This suggests that there is likely a specific context where expressing the *dma2* operon is advantageous. Even so, many factors can impact colonization fitness, and we only tested mice consuming a standard chow diet. Due to this, we cannot rule out that the ability of *Δdma2* to colonize *in vivo* may vary depending on the host diet and metabolic state. The gut microbiota also consists of many different bacterial species, so the lack of colonization fitness observed in *Δdma2* relative to the *Bt* WT begs the question of how this strain would compete in the presence of other *Bacteroides spp* and commensal microbes (**Fig. 7D, E**). Future studies are required to determine the exact role of Dma2, and other members of the Dma family, within the host.

## Materials and Methods

### Bacterial strains and Growth conditions

Strains, oligonucleotides, and plasmids are described in **Table S1** in the supplemental material. *Escherichia coli* was grown aerobically at 37°C in Luria-Bertani (LB) medium. *Bacteroides* strains were grown in an anaerobic chamber (Coy Laboratories) at 37°C containing an atmosphere of 10% H_2_, 5% CO_2_, 85% N_2_. *Bacteroides thetaiotaomicron* was cultured in Brain heart infusion (BHI) medium (Fisher Scientific) supplemented with 5 µg/ml Hemin and 1 µg/ml vitamin K3. When applicable, antibiotics were used as follows: 100 µg/ml ampicillin, 200 µg/ml gentamicin, 25 µg/ml erythromycin, and 10 μg/ml tetracycline. When required, Bacteroides was grown in minimal medium (MM) containing 100 mM KH2PO4 (pH 7.2), 15 mM NaCl, 8.5 mM (NH4)2SO4, 4 mM L-cysteine, 1.9 mM hematin/200 mM L-histidine (prepared together as a 1,000× solution), 100 mM MgCl2, 1.4 mM FeSO4.7H2O, 50 mM CaCl2, 1 µg/mL vitamin K3, and 5 ng/mL vitamin B12. Carbohydrates used to supplement MM include glucose, fructose, galactose, arabinose, mannose, rhamnose, xylose, amylopectin, gum arabic from Acacia Tree, levan, pectin from citrus peel, rhamnogalacturonan, *Saccharomyces cerevisiae* α-mannan, 1,6-β-glucan, heparin, hyaluronan, porcine mucin type II, porcine mucin type II, porcine mucin type III.

### Genetic Manipulation of *Bt*

We employed the pSIE1 vector described in Bencivenga-Barry et al. 2020 to develop constructs for generating clean deletion mutants in *Bt* (Bencivenga-Barry et al., 2020). Briefly, ∼750 base pair regions flanking our genes of interest were cloned into pSIE1. Completed vectors were then transformed into *E. coli s17λ-pir* to facilitate conjugation of the vector into *Bt*. Positive conjugants, containing the vector integrated into the *Bt* genome, were identified by selection on BHI plates containing gentamicin and erythromycin. To delete the gene of interest, counterselection was performed on BHI plates in the presence or absence of 125 ng/mL anhydrotetracycline (aTc). Mutants were identified by PCR prior to whole genome sequencing. Complemented strains were made using vectors from Whitaker et al. 2017 (Whitaker et al., 2017).

### OMV isolation

OMVs were purified by ultracentrifugation from cell-free culture supernatants according to our previously published methods (Elhenawy et al., 2014; Pardue et al., 2024; Sartorio et al., 2023; Valguarnera et al., 2018). Briefly, 50mL of *Bt* cultures grown to late stationary phase were centrifuged twice at 6,500 rpm at 4°C for 10 minutes. Supernatants were then filtered using a 0.22-µm-pore membrane (Millipore) to remove residual cells. The filtrate was subjected to ultracentrifugation at 200,000xg for 2 h (Optima L-100 XP ultracentrifuge; Beckman Coulter). Resulting supernatants were discarded, and the pellets, which contain OMVs, were resuspended in PBS standardized to OD_600_. When performing MS analysis, purified OMV preparations were lyophilized.

### Subcellular fractionation

<u>T</u>otal <u>M</u>embrane (TM) preparations were isolated by cell lysis and ultracentrifugation. Briefly, late stationary phase cultures were harvested by centrifugation at 6,500 rpm at 4°C for 10 minutes. The pellets were gently resuspended in a mixture of <u>P</u>hosphate <u>B</u>uffered <u>S</u>aline (PBS) containing complete EDTA-free protease inhibitor mixture (Roche Applied Science). Cells were then lysed using two passes through a cell disruptor at 35kPa. Next, centrifugation at 8,500 rpm at 4 °C for 8 minutes was performed to remove unbroken cells. Total membranes were collected by ultracentrifugation at 200,000xg for 1 h at 4°C. Supernatants were discarded, and pellets were resuspended in PBS standardized to OD_600_. TM fractions were lyophilized for MS analysis.

### SDS-PAGE and Western blot analyses

Total membrane and vesicle fractions were analyzed by standard 10% Tris-glycine SDS-PAGE. Samples were normalized by OD_600,_ and equivalent volumes were loaded onto an SDS-PAGE gel. Coomassie Blue staining was employed to analyze protein profiles. When applicable, samples were transferred onto a nitrocellulose membrane for Western blot analysis. Membranes were blocked using Tris-buffered saline (TBS)-based Odyssey blocking solution (LI-COR). Primary antibodies used in this study were rabbit polyclonal anti-His (ThermoFisher) and mouse monoclonal anti-FLAG (Sigma). Secondary antibodies used were IRDye anti-rabbit 780 (LI-COR). Imaging was performed using an Odyssey CLx scanner (LI-COR).

### Lipopolysaccharide (LPS) Silver stain

Abundance of LPS was measured according to Tsai and Frasch 1982 (Tsai & Frasch, 1982). Briefly, samples standardized by OD_600_, then diluted and treated with Proteinase K, before loading equal amounts onto a 15% SDS-PAGE gel. After running, gels were fixed overnight in 200 mL of 40% ethanol in 5% acetic acid. Next, gels were oxidized for 5min in 100mL of 0.7% fresh periodic acid in 40% ethanol and 5% acetic acid. Upon completion, gels underwent three washes (15 min each) in milliQ H_2_O. The gels were then stained for 10min in the dark with 28 mL 0.1M NaOH, 2mL NH_4_OH, 5mL 20% AgNO_3_, and 115 milliQ H_2_O. Gels underwent three additional washes prior to developing in 200 mL H_2_O with 10 mg Citric acid and 100 μL Formaldehyde.

### Growth assay with *Bt* strains

*Bt* WT and *Δdma2* strains were cultured overnight at 37 ⁰C under anaerobic conditions in BHI broth. Overnight cultures were centrifuged at 6500 rpm for 10 mins being washed with PBS. The cells were then resuspended in MM supplemented with the indicated carbon sources (0.5% w/v final concentration) and normalized to OD_600_ =0.1. Growth assays were conducted in sterile, round-bottom 96-well polystyrene microplates. Cultures were incubated anaerobically at 37 ⁰C under static conditions. OD_600_ readings were recorded every 30 mins using a Smart Reader 96-T (Accuris Instruments) following 10 seconds of orbital shaking to ensure homogenization. Each growth condition was tested in technical quadruplicate and independently repeated in three biological replicates.

### Cryo-ET sample preparation and imaging

*Bt* cells were grown on BHI-agar plates as described previously. Subsequently, cells were collected from the plate using a loop and resuspended in 1 mL of 1x PBS to a final OD_600_ of 3.0 for cryo-ET experiments. R2/2 carbon-coated 200 mesh copper Quantifoil grids (Quantifoil Micro Tools) were glow-discharged for 30 seconds. Then, cells were mixed with a solution of 10-nm gold beads treated with Bovine Serum Albumin. 4 μl of this mixture was applied on the grids in a Vitrobot chamber (FEI). Subsequently, the extra fluid was blotted off using a Whatman filter paper in the Vitrobot chamber with 100% humidity and the grids were plunge-frozen in a cryogen (liquid ethane). Samples imaging was performed at the Advanced Electron Microscopy at the University of Chicago. Cells were imaged using a Titan Krios transmission electron microscope operating at 300 kV and equipped with a BioQuantum K3 imaging filter (Gatan). Data were collected using Tomography 5 software with each tilt series ranging from -60° to 60° in 3° increments with a pixel size of 3.35 Å, an underfocus of 7 μm and a total dose of 130 e^-^/Å^2^. Subsequently, three-dimensional reconstructions of tilt-series and further visualization were performed using the IMOD software package (Kremer et al., 1996).

### RNA Sequencing Sample Collection, Library Preparation, and Analysis

RNA was isolated from *Bt* cells according to our methods outlined in Pardue et al. 2024 (Pardue et al., 2024). Briefly, WT and *Δdma2* were grown overnight in BHI media before being diluted to the equivalent of OD 0.1 in 10 mL and grown anaerobically for 4 h at 37°C. Four individual overnight and 10-mL culture biological replicates were prepared. Cultures were normalized, and an amount of culture equivalent to an OD_600_ of 4.0 was pelleted for 90s at 8,000 rpm. Pellets were resuspended on ice in 1 mL TRIzol (Invitrogen) with 10 μL of 5 mg/mL glycogen. Samples were flash frozen and stored at −80°C until extraction. Prior to extraction, samples were thawed on ice, then pelleted, and supernatants were treated with chloroform. RNA was extracted from the aqueous phase using the RNeasy minikit (Qiagen, Inc.), and RNA quality was checked by agarose gel electrophoresis and *A*_260_/*A*_280_ measurements. RNA was stored at −80°C with SUPERase-IN RNase inhibitor (Life Technologies) until library preparation.

RNA sequencing prep (RNA-Seq) was performed as previously describe (Zhu et al., 2020). Briefly, 400ng of total RNA from each sample was used for generating cDNA libraries following our RNAtag-Seq protocol. PCR amplified cDNA libraries were sequenced on an Illumina NextSeq500, obtaining a high-sequencing depth (over 7 million reads per sample). RNA-Seq data was analyzed using our *in-house* developed analysis pipeline *Aerobio*. Raw reads are demultiplexed by 5’ and 3’ indices, trimmed to 59 base pairs, and quality filtered (96% sequence quality>Q14). Filtered reads are mapped to the corresponding reference genomes using bowtie2 with the --very-sensitive option (-D 20 –R 3 –N 0 –L 20 –i S, 1, 0.50). Mapped reads are aggregated by feature Count and differential expression is calculated with DESeq2 (Zhu et al., 2020). In each pair-wise differential expression comparison, significant differential expression is filtered based on two criteria: |log2foldchange| > 1 and adjusted p-value (*padj*) <0.05. All differential expression (DE) comparisons are made between the WT and ΔDma1 mutants under the condition mentioned above. The reproducibility of the transcriptomic data was confirmed by an overall high Spearman correlation across biological replicates (R > 0.95). BioProject: PRJNA1298834.

### Sample preparation for Proteomic analysis

WT, *Δdma2*, and *Δdma2-das2* were grown overnight anaerobically in 3mL of BHI media prior to being diluted into 50 mL and grown for 20 h. Whole cells, total membranes, and vesicles were collected from each strain. Four individual biological replicates of each fraction were performed for each strain. Samples were lyophilized in preparation for MS analysis.

### Proteomic analysis

Acetone precipitated protein biological replicates/fractions were solubilized in 4% SDS, 100 mM HEPES by boiling for 10 min at 95 °C, then protein concentrations assessed using bicinchoninic acid protein assays (Thermo Fisher Scientific). 200 μg of each biological replicate/fraction was prepared for digestion using S-trap mini columns (Protifi, USA) according to the manufacturer’s instructions. Briefly, samples were reduced with 10 mM dithiothreitol for 10 mins at 95 °C and then alkylated with 40 mM Iodoacetamide in the dark for 1 hour. Samples were acidified to 1.2% phosphoric acid and diluted with seven volumes of S-trap wash buffer (90% methanol, 100 mM Tetraethylammonium bromide pH 7.1) before being loaded onto S-traps and washed 3 times with 400 μL of S-trap wash buffer. Samples were then digested with 4 μg of Trypsin (a 1:50 protease/protein ratio) in 100 mM Tetraethylammonium bromide overnight at 37 °C before being collected by centrifugation with washes of 100 mM Tetraethylammonium bromide, followed by 0.2% formic acid, and then 0.2% formic acid / 50% acetonitrile. Samples were dried down and further cleaned up using C18 Stage ^1,2^ tips to ensure the removal of any particulate matter.

C18 cleaned up peptide samples were re-suspended in Buffer A* (2% acetonitrile, 0.1% trifluoroacetic acid in Milli-Q water) and separated using a two-column chromatography set-up on a Dionex Ultimate 3000 UPLC composed of a PepMap100 C18 20 mm x 75 μm trap and a PepMap C18 500 mm x 75 μm analytical column (Thermo Fisher Scientific) coupled to a Orbitrap Fusion™ Lumos™ Tribrid™ Mass Spectrometer (Thermo Fisher Scientific) with a FAIMS Pro interface (Thermo Fisher Scientific). 145-minute gradients were run for each sample, with samples loaded onto the trap column with 98% Buffer A (2% acetonitrile, 0.1% formic acid in Milli-Q water) and 2% Buffer B (80% acetonitrile, 0.1% formic acid) with peptides separated by altering the buffer composition from 2% Buffer B to 28% B over 126 minutes, then from 28% B to 40% B over 9 minutes, then from 40% B to 80% B over 3 minutes, the composition was held at 80% B for 2 minutes, and then dropped to 2% B over 2 minutes and held at 2% B for another 3 minutes. A data-dependent stepped FAIMS approach was utilised with two different FAIMS CVs of -45 and -65, as previously described ^3^. For each FAIMS CV a single Orbitrap MS scan (500-2000 m/z, maximal injection time of 50 ms, an AGC of maximum of 4*10^5^ ions and a resolution of 60k) was acquired every 2 seconds followed by Orbitrap MS/MS HCD scans of precursors (NCE 30%, maximal injection time of 80 ms, an AGC set to a maximum of 1.25*10^5^ ions and a resolution of 30k).

### Proteomic data analysis

Prior to Identification and LFQ analysis files were separated into individual FAIMS fractions using the FAIMS MzXML Generator ^4^. Separated FAIMS fractions were searched against the *Bacteroides thetaiotaomicron* VPI-5482 proteome (Uniprot: UP000001414) using MaxQuant (v1.6.17.0)^5^ allowing carbamidomethylation of cysteine set as a fixed modification and oxidation of methionine as a variable modification. Searches were performed with Trypsin cleavage specificity, allowing 2 miscleavage events with a maximum false discovery rate (FDR) of 1.0 % set for protein and peptide identifications. The LFQ and “Match Between Run” options were enabled to allow comparison between samples. The resulting data files were processed using Perseus (v1.4.0.6)^6^ with missing values imputed based on the total observed protein intensities with a range of 0.3 σ and a downshift of 1.8 σ. Statistical analysis was undertaken in Perseus using two-tailed unpaired T-tests.

### Data availability

The mass spectrometry proteomics data has been deposited in the Proteome Xchange Consortium via the PRIDE partner repository with the data set identifier: PXD066605 Username: Password: D8Oefcsz1pfv.

### Negative staining and analysis by transmission electron microscopy

For quantitative analyses at the ultrastructural level, 200 mesh formvar/carbon-coated copper grids (Ted Pella Inc., Redding CA) were coated with 50µg/ml poly-L-lysine (Sigma, St Louis, MO) for 10 min at 37^○^C. Excess fluid was removed, and grids were allowed to air dry. Poly-L-lysine coating allowed for even distribution of material across the grid. Bacterial OMVs were fixed with 1% glutaraldehyde (Ted Pella Inc.) and allowed to absorb onto freshly glow discharged poly-L-lysine-coated grids for 10 min. Grids were then washed in dH_2_O and stained with 1% aqueous uranyl acetate (Ted Pella Inc.) for 1 min. Excess liquid was gently wicked off and grids were allowed to air dry. Samples were viewed on a JEOL 1200EX transmission electron microscope (JEOL USA, Peabody, MA) equipped with an AMT 8-megapixel digital camera (Advanced Microscopy Techniques, Woburn, MA). Each OMV sample was processed in triplicate (3 grids). Ten random images were taken at a magnification of 25,000x from various areas of the grid with a total of 90 images for each sample, and the number of OMV in each image was quantified.

### Competitive colonization of Antibiotic-treated mice

All animal experiments were approved by the Washington University Animal Care and Use Committee, and we have complied with all relevant ethical regulations. All mice used were from the inbred C57/BL6 line. Six-week-old animals were used for colonization experiments. Mice were administered an antibiotic cocktail consisting of ampicillin (333.3 mg/ml), neomycin (333.3 mg/ml), metronidazole (10 mg/ml), and vancomycin (166.7 mg/ml), each mouse receiving 150μl by oral gavage, every 24 hrs for 7 consecutive days to deplete the normal intestinal flora. Next, mice were given an inoculum of a single *Bt* strain, for monocolonization experiments, or two *Bt* strains, for co-colonization experiments (∼10^10^ CFUs/oral gavage total; an aliquot was taken from the input inoculum and plated on BHI agar to count CFUs) for 2 consecutive days. Fresh fecal pellets were collected 2, 3, 5, 7, 10, and 14 days post oral gavage and used to quantify CFU/ml to track colonization throughout the duration of the experiment. Four mice were utilized per condition, and each experiment was conducted in triplicate for a total of twelve mice per condition. Competitive index represents the ratio of mutant CFU/g of feces to that of the WT.

## Acknowledgements

This work was supported by funding M.F.F. (R01AI181213) through the National Institute of Allergy and Infectious Diseases of the National Institutes of Health. N.E.S was supported by an Australian Research Council Future Fellowship (FT200100270), an ARC Discovery Project Grant (DP210100362) and a NHMRC Ideas Grant (GNT2018980). We thank the Melbourne Mass Spectrometry and Proteomics Facility of The Bio21 Molecular Science and Biotechnology Institute for access to MS instrumentation.

---

**Supplemental Figure 1:**
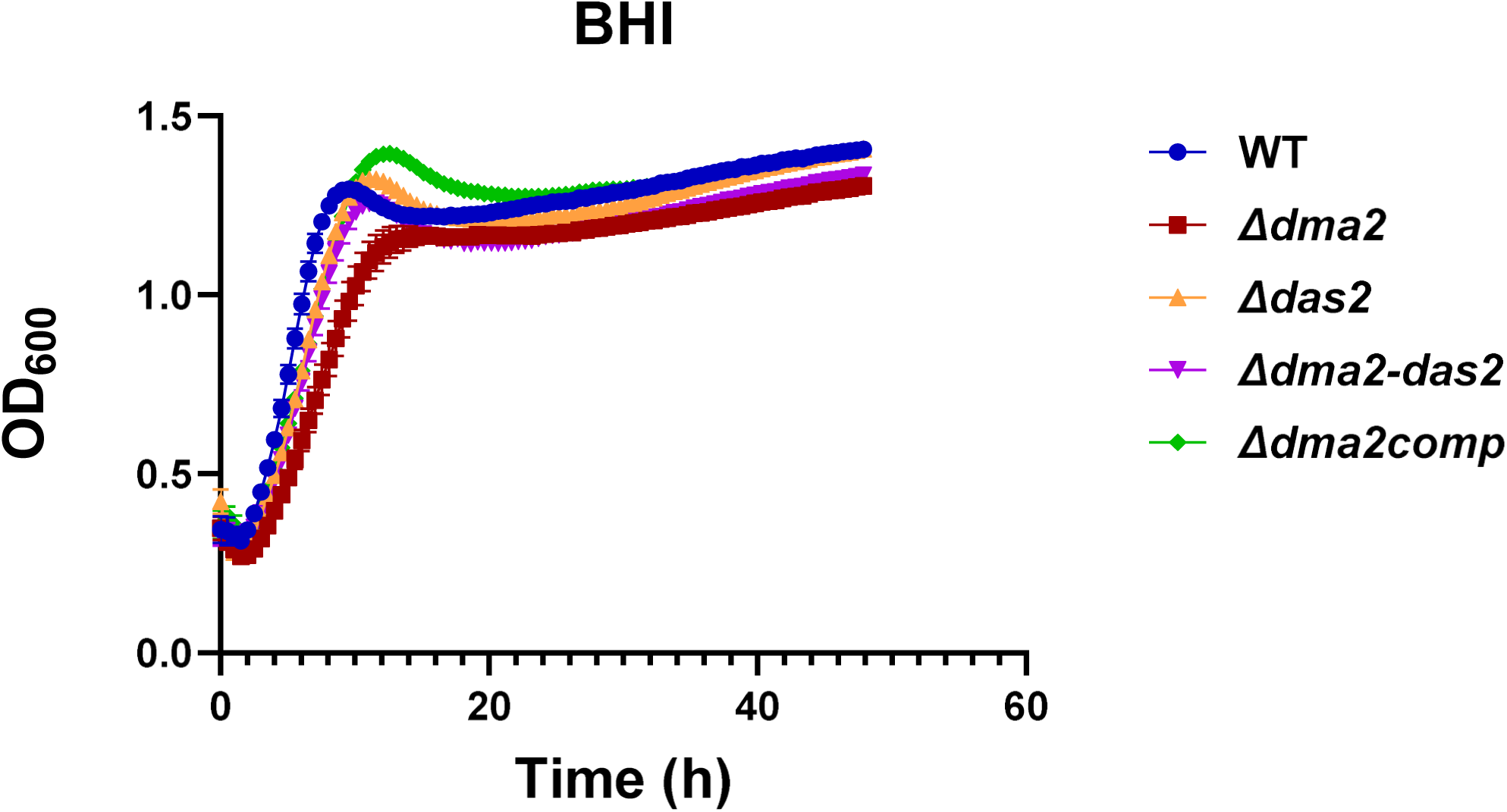
Mutation of *das2* does not impact growth *in vitro*. Growth curves showing the growth of *Bt* WT, *Δdma2*, *Δdas2*, *Δdma2-das2*, and *Δdma2_Comp_* in supplemented BHI broth. Growth curves presented are the average from the results of three independent biological replicates each including four technical replicates from each strain.

**Supplemental Figure 2:**
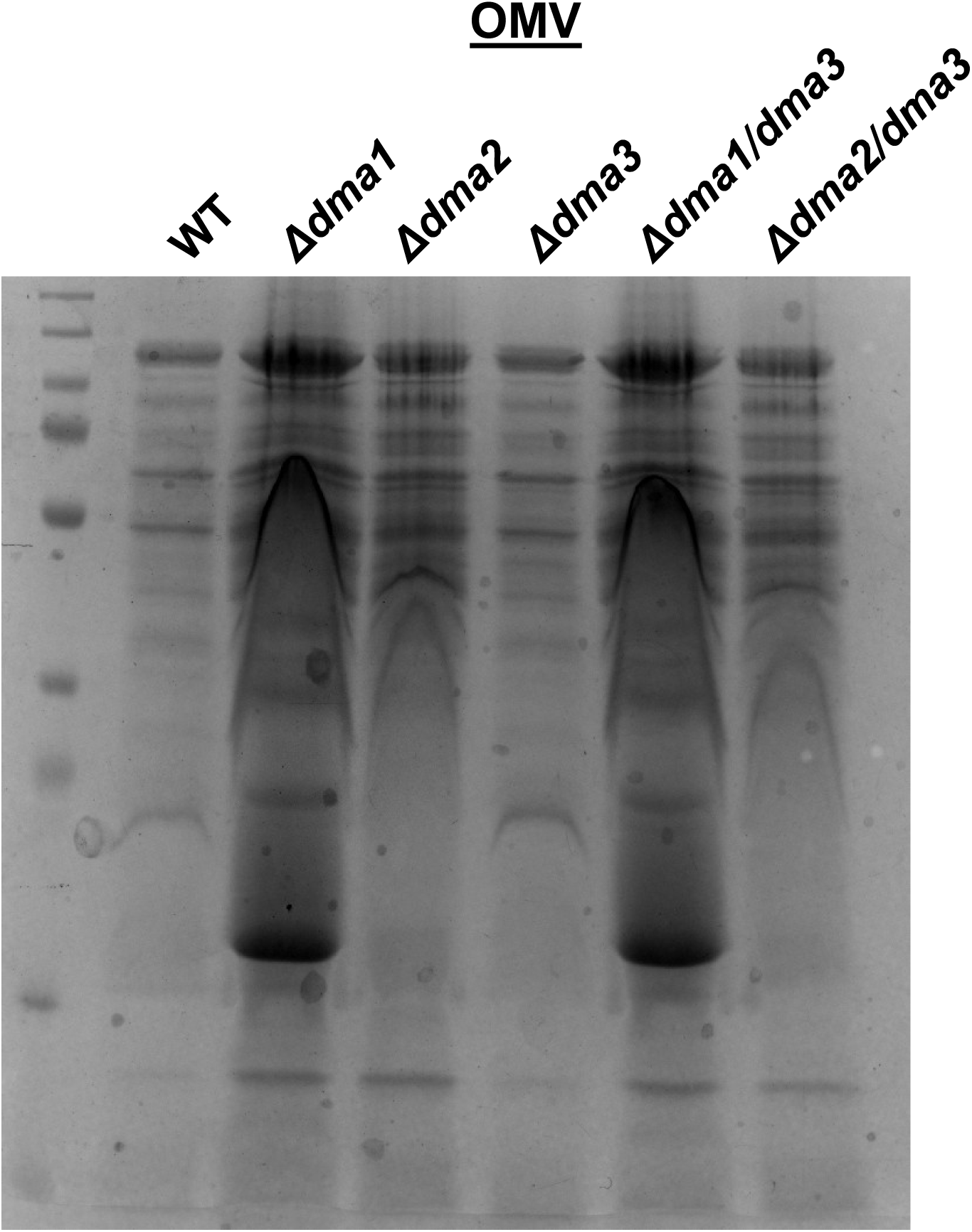
Dma3 does not promote OMV biogenesis in *Δdma1* and *Δdma2.* Coomassie Blue stain of OMV fractions isolated from *Bt* WT, *Δdma1, Δdma2*, and the corresponding strains containing deletion in *dma3*. Samples were normalized by OD_600_ and analyzed by 10% SDS-PAGE. This shows that mutation of *dma3* does not impact the electrophoretic profile of the OMV fraction in any of the genetic backgrounds tested.

**Supplemental Figure 3:**
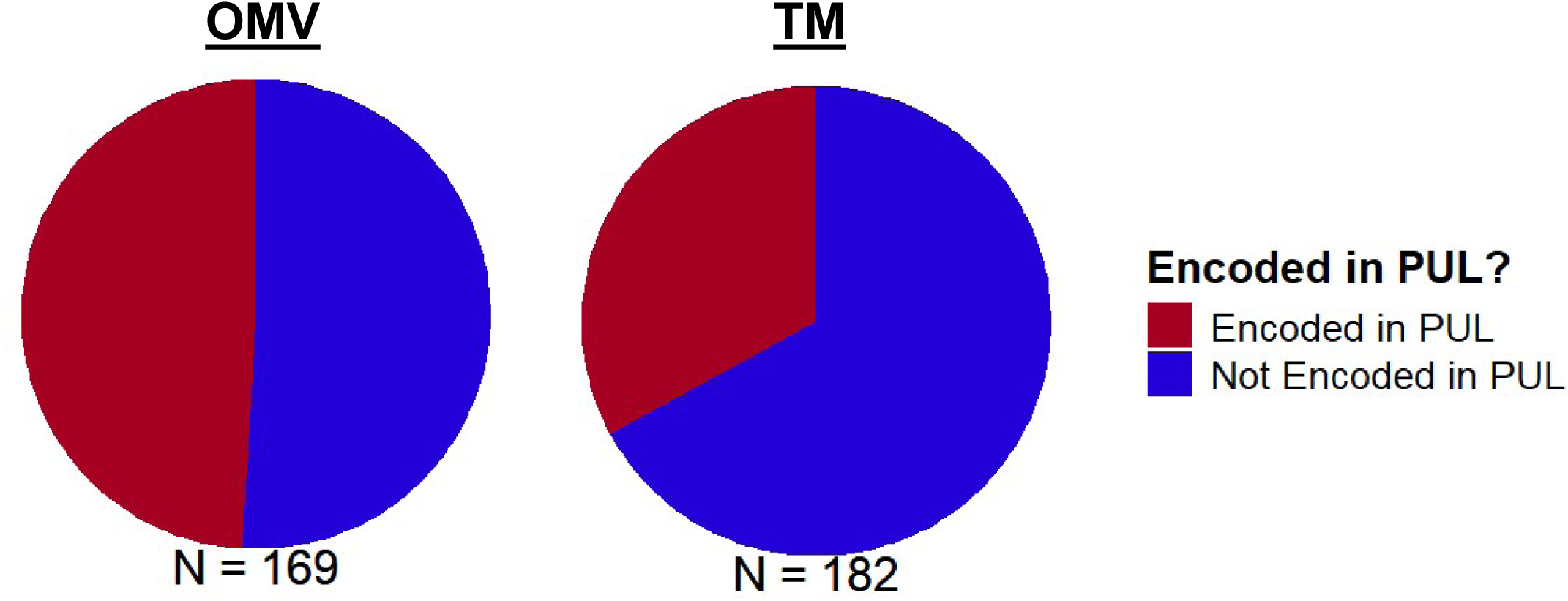
PUL encoded proteins are the main group of proteins altered in the OMV fraction of *Δdma2*. Pi charts corresponding to the proportion of significantly altered proteins from the *Bt Δdma2* and *Δdma2-das2* proteomics encoded in PULs **(Dataset S6, S7).**

**Supplemental Figure 4:**
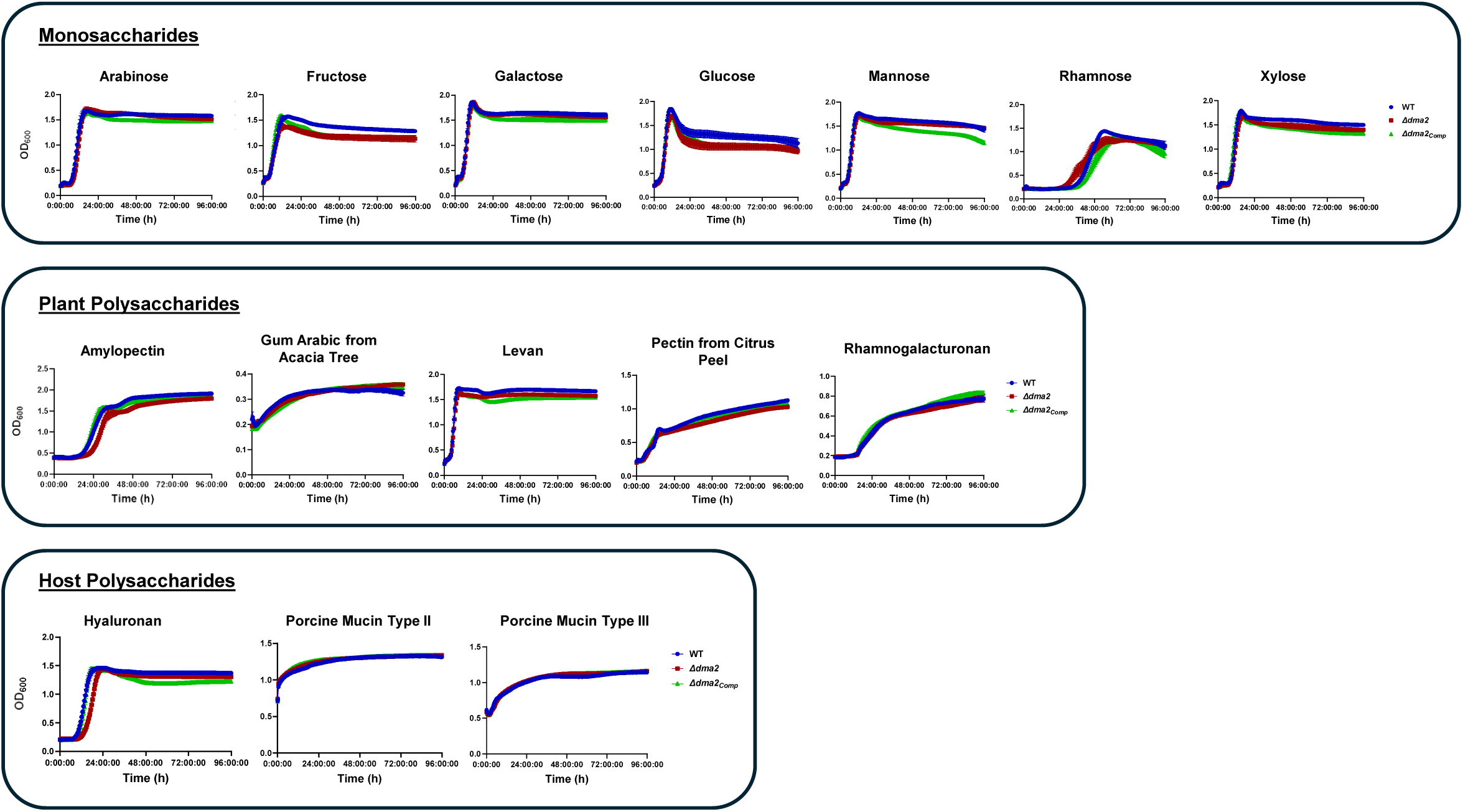
Growth of *Δdma2* in the presence of various carbon sources. Growth curves showing the growth of *Bt* WT, *Δdma2*, and *Δdma2_Comp_* in the presence of minimal media containing various monosaccharides and polysaccharides. Growth curves were generated from the results of three independent experiments each including four technical replicates from each strain

**Supplemental Figure 5:**
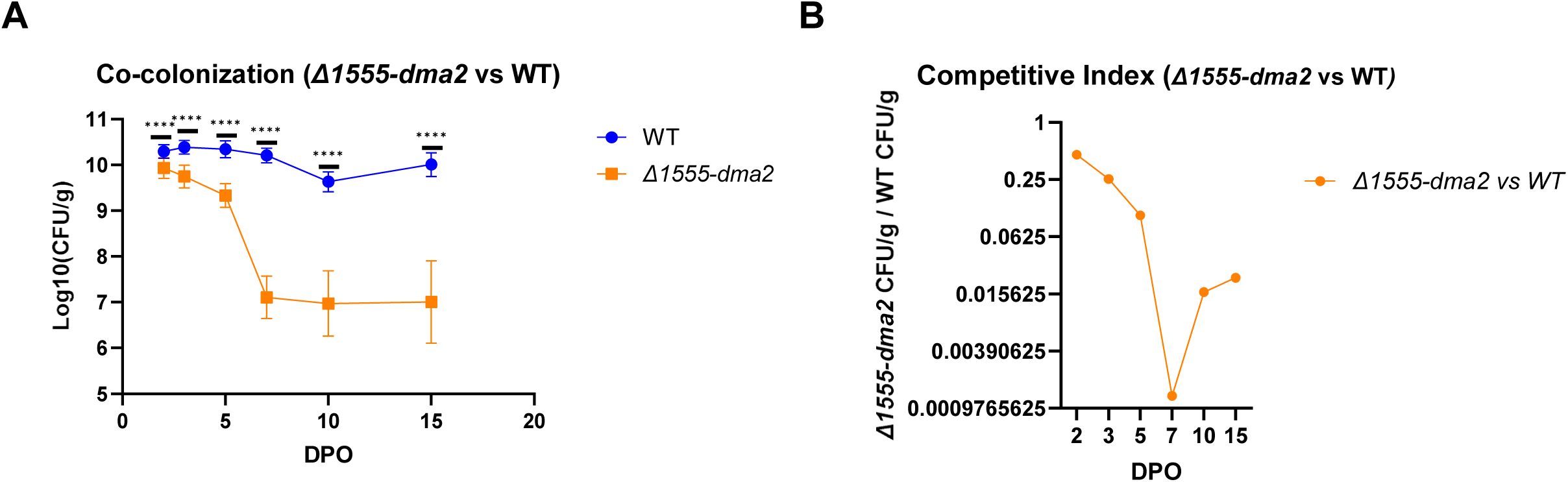
Fitness defect *in vivo* is not due to the overexpression of *dma2* operon in *Δdma2*. (A, B) Co-colonization experiment comparing *Bt* WT and *Δ1555-dma2*. This shows that *Δ1555-dma2* experiences and immediate decrease in abundance when the WT is present. This phenotype is similar the that observed in *Δdma2*, which indicates that the overexpression of the downstream genes in the *dma2* operon is not what causes the decline in colonization fitness observed in *Δdma2*. Points on the graph represent the mean and standard deviation of data collected from three independent experiments containing four mice each per condition. Significance threshold corresponds to: (*) p-value = 0.05, (**) p-value = 0.01, (***) p-value = 0.001, (****) p-value = 0.0001.

**Supplemental Dataset 1: RNA Sequencing-*Δdma2* vs *WT***

**Supplemental Dataset 2: Polysaccharide Utilization Loci in *Bacteroides thetaiotaomicron***

**Supplemental Dataset 3: Genes Upregulated in *Δdma2* and *Δdma1***

**Supplemental Dataset 4: Genes Downregulated in *Δdma2* and *Δdma1***

**Supplemental Dataset 5: RNA Sequencing-*Δdma2* vs *Δdma2-das2***

**Supplemental Dataset 6: OMV Proteomics-*Δdma2* vs *Δdma2-das2***

**Supplemental Dataset 7: TM Proteomics-*Δdma2* vs *Δdma2-das2***

